# Maximizing the coding capacity of neuronal networks

**DOI:** 10.1101/673632

**Authors:** Sandeep Chowdhary, Collins Assisi

## Abstract

Information in neuronal networks is encoded as spatiotemporal patterns of activity. The capacity of a network may thus be thought of as the number of stable spatiotemporal patterns it can generate. To understand what structural attributes of a network enable it to generate a profusion of stable patterns, we simulated an array of 9 *×* 9 neurons modelled as pulse-coupled oscillators. The structure of the network was inspired by the popular puzzle Sudoku such that its periodic responses mapped to solutions of the puzzle. Given that there are nearly a 10^9^ possible Sudokus, this networks could possibly generate 10^9^ spatiotemporal patterns. We show that the number of stable patterns were maximized when excitatory and inhibitory inputs to each neuron were balanced. When this balance was disrupted, only a subset of patterns with certain symmetries survived.

---

The sense of smell [1], neural representations of movement [2], and episodic memories [3] are all instantiated in the wetware of the brain as the activity of interconnected networks of neurons. The connections between neurons can be excitatory or inhibitory. Synaptic inhibition, where the activity of a neuron tends to suppress the activity of others it connects, is central to orchestrating these patterns of activity [4]. In an inhibitory network, when a group of neurons that spike in synchrony, stop firing, it releases others from inhibition. These, in turn, start firing before they yield to another group. This pattern of activation in competitive networks has been termed winnerless competition [5]. The identity of synchronously spiking neurons and the order in which they fire are determined by the structure of the network [6]. Competitive interactions between neurons may be usefully related to the vertex coloring [7] of the coupling network [8, 9]. Neurons (vertices) associated with different colors inhibit each other, causing them to spike at different times while those associated with the same colour can fire at the same time. Therefore, a spatiotemporal pattern of activity may be mapped to a coloring of the network. There are typically a large number of ways to color a random network. This explosion of possibilities also implies that an inhibitory network can potentially generate a large number of spatiotemporal patterns. These patterns may be thought of as neural representations of odors, sequences of movement or episodic memories. The questions, how many patterns can a network elicit (the capacity of the network), how stable are these patterns and can the network flexibly switch between different patterned outputs, thus become central to understanding brain function. To address these questions we sought to construct networks that were sufficiently complex, in that they possessed many colorings (equivalently, allowed sequences of activity). However, finding all colorings [10] and simulating the dynamics of a large random network is a formidable problem. As with many complex systems, the key often lies in identifying a system that is simple enough to work with, but sufficiently complex that it embodies many of the essential components of the original, seemingly intractable system. We discovered that the simple and popular puzzle, Sudoku, provides an appropriate framework to address our questions.

In this paper, we examine the relationship between the topology of a network of pulse coupled oscillators and the dynamics of its constituent elements in the context of a model network inspired by Sudoku. A Sudoku puzzle consists of a 9 × 9 grid. Integers from 1 to 9, termed clues, are placed in different cells of the grid. The objective of the puzzle is to fill in the remaining cells such that each integer occurs only once in each row, column and 3 × 3 sub-grid. Typically a unique combination of numbers solves the puzzle for a given set of clues. As one removes clues from the puzzle, the number of possible solutions increase. In the absence of any clues, the total number of possible ways in which one can populate the grid while satisfying the constraints is 6, 670, 903, 752, 021, 072, 936, 960. When solutions that share certain symmetries are eliminated this number comes down to 5, 472, 730, 538 [11]. Each of these solutions is a 9−coloring of the network illustrated in The in the same row, column and sub-grid. Here we examine the dynamics of a network of neurons connected according the topology described above. The neurons that occupied the vertices of the graph were modeled as a Mirollo-Strogatz pulse-coupled oscillators [12], a generalization of the integrate and fire neuron [13], given by the equation,

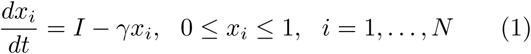

where, when *x*_*i*_ reached a threshold (*x*_*thresh*_ = 1), the oscillator emitted a pulse and was reset to *x*_i_ = 0. The system oscillated with a period 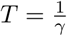 in 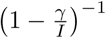. The solution of equation(1) is

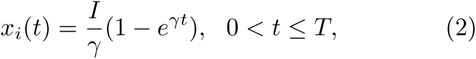

*x* can be expressed as a function, *U,* of a phase varible *φ*, where *U*: [0, 1] *→* [0 1] is a monotonically increasing (*U′* > 0) and concave down (*U″* < 0) function. The phase of the oscillator increased monotonically according to the equation 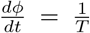. When it reached a threshold, *φ* _*thresh*_ = 1, it emitted a pulse and was re-set to *φ* = 0. A pulsatile interaction between oscillators advanced or retarded the phase of the oscillators it connected. The extent to which the phase of the receiving oscillator shifted was determined by the function *U* (*φ*). By setting 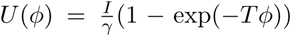, we have *U* (*φ*).*=x(φT/γ)* Thus, each oscillator behaved as an integrate-and-fire neuron with period *T*. Excitatory inputs to an oscillator advanced the phase as follows,

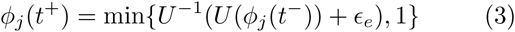

where *ϵ* _*e*_ = Θ(*s*) is the Heaviside step function and *s* is the number of pulses received at a given time. Inhibitory inputs, in contrast, retard the phase of the receiving oscillator.

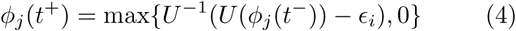

Excitatory interactions synchronize the oscillators (1). In two reciprocally coupled oscillators each excitatory pulse advanced the phase of the receiving oscillator towards the threshold until the spike times of both eventually coincided (Figure 2a,b). This form of synchrony is seen even in larger all-all coupled networks [12]. We assume here that interaction delays are negligible. When delays were present, purely excitatory networks of pulse-coupled oscillators generate elaborate spatiotemporal patterns of spikes [14]. A pair of inhibitory oscillators, in contrast, spike at different phases that progressively separate over multiple cycles (Figure 2b,bottom traces).

**FIG. 1:**
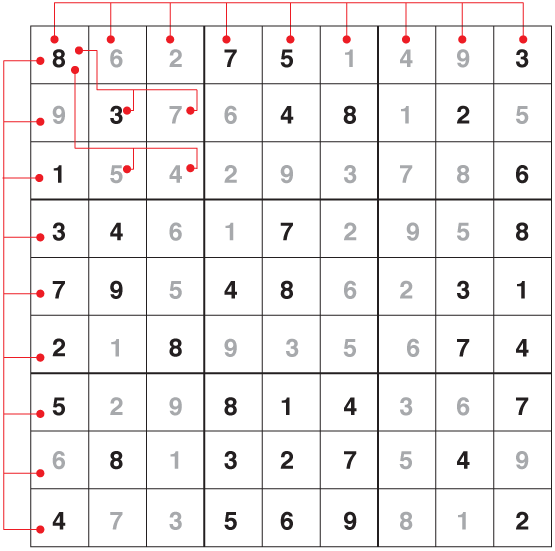
A Sudoku puzzle. The numbers in the dark font are the ‘clues’. To find a solution, one fills in the remaining squares with numbers (rendered in gray) such that every number from 1 to 9 occurs once in each row, column and subgrid (separated by heavier lines). Every Sudoku solution is a 9 −coloring of the network shown in red. The lines indicate the connections to and from the top left element - 8. All other nodes are connected in a similar manner. The solution shown here appeared in the the World Sudoku championship, March 2006.

**FIG. 2:**
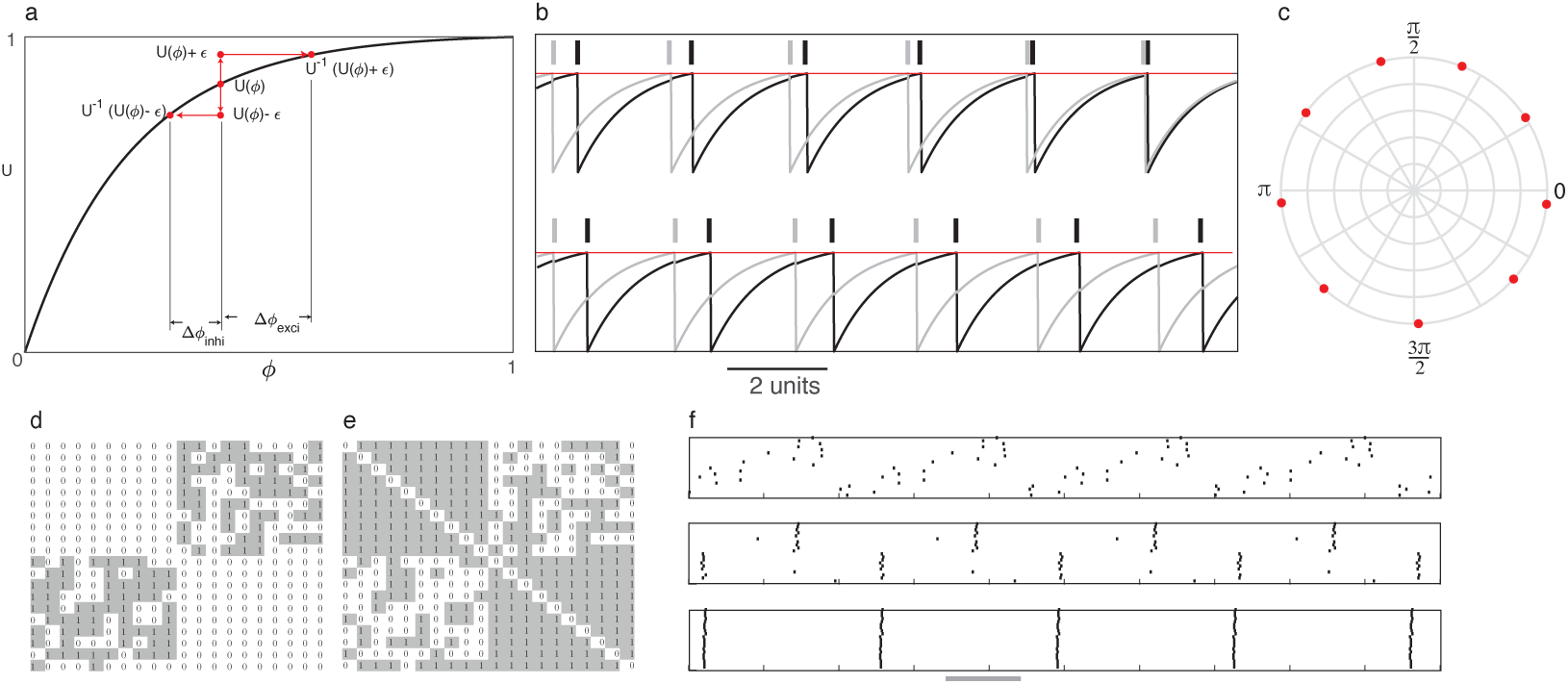
**(a)**The function *U* (*φ*). An excitatory pulse advances the phase by Δ*φexci*. An inhibitory pulse retards the phase by Δ*φ*_*inhi*_ **(b)** Dynamics of two oscillators coupled by reciprocal excitation(top traces) and inhibition (bottom traces). Lines show the time when pulses are emitted as *x* crosses the threshold (red line) **(c)** Splay phase state of a network of 9 all-all coupled oscillators **d,e** Adjacency matrices of the inhibitory and excitatory networks. **(f)**Response of the bipartite network when excitation was low (top panel), balanced with inhibition (middle panel) and high (bottom panel)

**FIG. 3:**
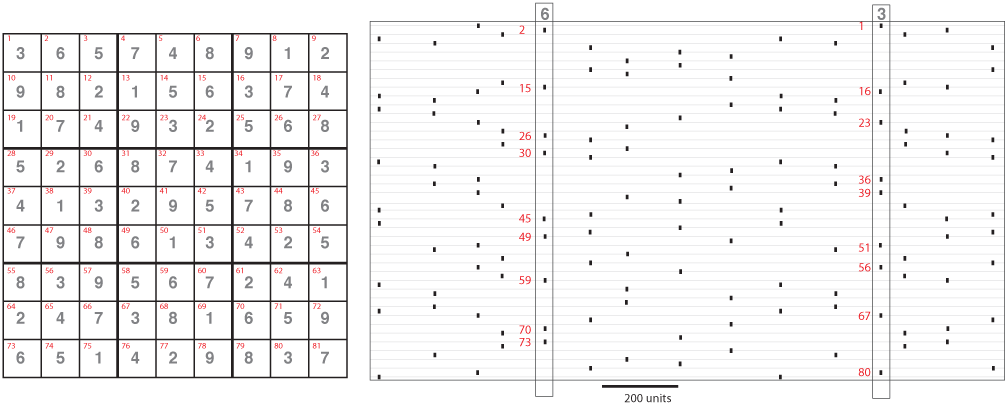
Mapping a periodic attractor to a Sudoku puzzle solution. The oscillators (indexed in red) that spiked within a short time window (marked in the right plot). The indices matched that of the Sudoku solution (Matrix on left).

How does the dynamics map to vertex colorings of the coupling graph? Consider a graph *G* = (*V, E*) where *V* are the vertices and *E* ⊆ [*V]*^2^ are the edges of *G*. A vertex coloring of *G* is a mapping *C*: *V → S* such that Figure(1) where each node is connected to every node *C*(*v*) *≠ C*(*w*) whenever *v* and *w* are adjacent [15]. elements of *S* are called colors. We will be interested in minimal colorings where the number of elements *S* = *k* is the least required to color *G*. Now, consider an all-all connected network of 9 oscillators. Since *C*(*v*) ≠ *C*(*w*) for all pairs of vertices, each has to be assigned a different color, that is, *S* has 9 elements, say *S* = {1, 2*, …* 9}. We simulated this network and calculated the phase of all the oscillators every time one of the oscillators (say 1), fired a spike (*φ* = 0). The phases the oscillators were distributed between 0 and 2*π* (Figure 2). An oscillator with a particular phase could thus be mapped to an element of *S* are called colors. We will be interested in minimal colorings where the number of elements *S* = *k* is the least required to color *G*. Now, consider an all-all connected network of 9 oscillators. Since *C*(*v*) = *C*(*w*) for all pairs of vertices, each has to be assigned a different color, that is, *S* has 9 elements, say *S* = 1, 2*, …* 9. We simulated this network and calculated the phase of all the oscillators every time one of the oscillators (say 1), fired a spike (*φ* = 0). The phases the oscillators were distributed between 0 and 2*π* (Figure 2). An oscillator with a particular phase could thus be mapped to an element of *S* are called colors. We will be interested in minimal colorings where the number of elements *S* = *k* is the least required to color *G*. Now, consider an all-all connected network of 9 oscillators. Since *C*(*v*) = *C*(*w*) for all pairs of vertices, each has to be assigned a different color, that is, *S* has 9 elements, say *S* = 1, 2*, …* 9. We simulated this network and calculated the phase of all the oscillators every time one of the oscillators (say 1), fired a spike (*φ* = 0). The phases the oscillators were distributed between 0 and 2*π* (Figure 2). An oscillator with a particular phase could thus be mapped to an element of *S*. Does this mapping between the phase dynamics and the vertex colorings extend to arbitrary networks? To test this we first simulated a bipartite network consisting of two groups that inhibited each other. The probability of connections between groups was set to 0.6. We did not implement any within-group connections in this network (Figure 2d). For the bipartite network simulated in Figure (2) a minimum of two colors were required to partition the graph and only one 2 coloring was possible. Our simulations showed that while connected neurons did not fire together, nodes from the same partition did not fire synchronously either. In random bipartite networks, each oscillator from a partition received inhibitory inputs from some randomly chosen sub-group of the oscillators of the other partition. Oscillators within a partition did not synchronize since they received heterogeneous inputs (Figure 2, top panel). Since excitatory interactions were absent, all oscillators from the same partition were not drawn to synchronize. Here, the dynamics did not always map to a minimal coloring of the coupling network. Next, we constructed a complementary excitatory network viz. we connected an excitatory edge between nodes that did not inhibit each other (see figure 2d,e) for the respective adjacency matrices). This network (Figure 2e) synchronized oscillators within each group. However anti-phase synchrony between groups was possible only within a regime where the cumulative strength of excitatory input was balanced by inhibitory inputs(Figure 2f, middle panel). When the ratio of excitatory to inhibitory input was large the excitatory drive within and across groups drove all the oscillators, even those belonging to different partitions, to spike together (Figure 2f, bottom panel).

Unlike the bipartite graph simulated here, in general a network may possess multiple colorings. For example, the Sudoku network 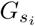 Figure(1), where the vertices along a row, column and sub-grid are all-all connected, may be colored in *∼*10^9^ different ways. We constructed a balanced Sudoku network where inhibitory interactions between nodes were given by 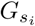 (see Figure 1) and excitatory interactions were given by the complementary network 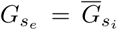 The network did not possess any self-loops. Our simulations showed that after a transient, the system settled to periodic attractors where different groups of oscillators spiked sequentially. The transient time to arrive at these attractors varied widely for different initial conditions and excitatory/inhibitory ratios. To determine whether the observed periodic attractors could be mapped to 9 −colorings of 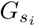(valid Sudoku solutions), we assigned the same number to oscillators that spiked together. For example, the number 3 was located at the indices 1, 16, 23*, …,* 76 of the matrix and matched the identity of neurons that spiked at a particular time(see figure(3)). Similarly, we matched the other cells of the Sudoku grid by associating numbers to synchronously spiking oscillators until all the cells were populated. We could always find a minimal coloring of 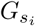 within one cycle of the periodic attractor orbit. Different colorings were associated with different clusterings of the phases of the 81 pulse-coupled oscillators.

The coloring of the network can be used to identify which groups of oscillators spike together. Here we ask, for a given coloring, can one determine the phase ordering of the oscillators around the circle. In a complete −9 partite inhibitory network, each node in a cluster received inputs from all the neurons in the remaining 8 clusters. There were no within-group connections. The symmetry of the network was evident in the dynamics of the oscillators. They formed 9 clusters that spiked periodically in succession. The phase difference between neighboring clusters (alternately the time between spikes), was nearly identical. Any permutation of the phase ordering of the synchronous firing groups was possible (Figure 5b). Therefore, there were 8! possible sequences that this network could generate. Solutions to Sudoku puzzles are −9 colorings of 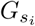, but do not have the same symmetries as the coloring of a complete 9 − partite network. To understand how the symmetries of a particular coloring constrain the phases of the oscillators, we defined a matrix *J* (*C*), for each coloring *C*. The *ij*^*th*^ element of *J* (*C*) was the number of inhibitory connections between the two groups of oscillators, each group being of a particular color. The groups were labeled *P*_*i*_ and the *P*_*j*_ for a particular coloring *C* of the network 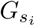, A vertex in a partition, say *P*_*i*_, is connected to either two or three vertices from any other partition *P*_*j*≠*i*_ (figure 4a). For example, the vertex labeled 7 in figure 4a is connected to two vertices labeled 3 and 3 vertices labeled 9. Suppose a vertex is connected to 2 vertices in *N*_2_ out of 8 partitions and to 3 vertices in *N*_3_ partitions. 2*N*_2_ + 3*N*_3_ = 20 since the degree of every vertex in the network is 20. The total number of partitions a vertex in any partition *P* can connect to is *N*_2_ +*N*_3_ = 8. Therefore, *N*_2_ = *N*_3_ = 4. That is, every vertex has 2 neighbors in exactly 4 partitions and 3 neighbors in the remaining 4 partitions. Therefore, the elements of *J* (*C*), viz. the number of inhibitory connections between partitions, lies between 18 and 27 for any 9−coloring *C*.

**FIG. 4:**
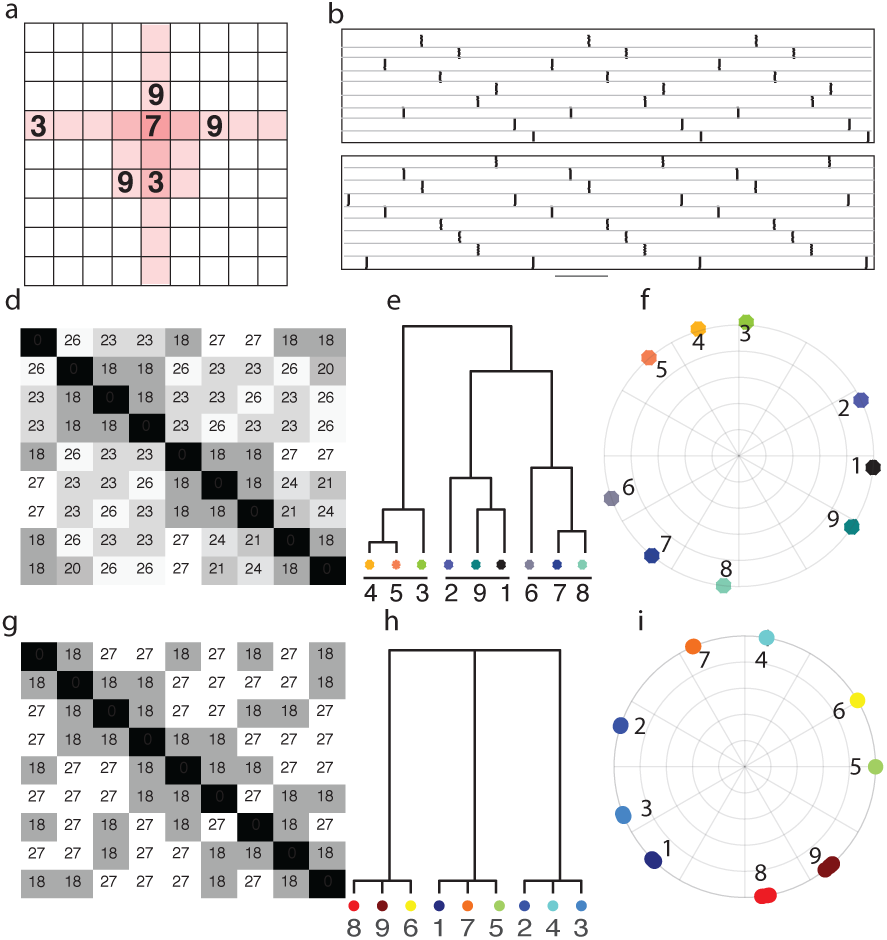
**(a)** Each vertex connects to either 2 or 3 vertices of a different color. 7 connects 2 vertices marked 3 an 3 vertices marked 9**(b)**Dynamics of a complete 9 − partite network. Each dot marks the location of a spike that the neuron (indexed along the y-axis) fired. 9 synchronously spiking groups of 9 neurons are shown for two intial conditions (top and bottom panel) **d,g***J* (*C*) matrices for two colorings. **e,h** Hierarchical clustering based on pairwise distances between clusters calculated from *J* (*C*). **e,f** Different splay phase states of the network in Figure(1)

**FIG. 5:**
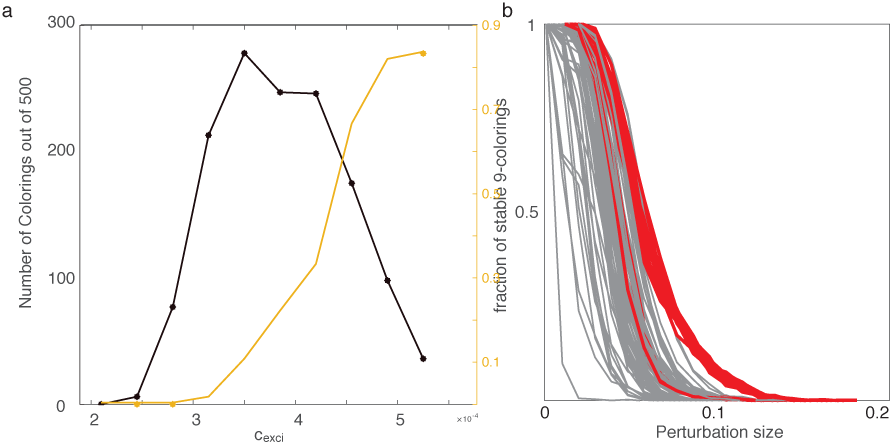
**(a)**The number of unique periodic attractors plotted as a function of the strength of excitation. Also shown is the fraction of 18 − 27 attractors (yellow trace). **(b)**Each line shows the proportion of solutions (out of 500) that persist as the strength of the perturbation increases. The red lines show the proportion for 18 − 27 colorings. All other colorings are shown in gray.

Larger *J*_*ij*_(*C*) meant that the groups *i* and *j* collectively inhibited each other to a greater extent than groups with fewer inhibitory connections. Thus we expect the phase difference between these groups of oscillators to be correspondingly larger. We used the matrix *J* (*C*) as a measure of the pairwise distance between the 9 clusters. The distance between groups was used to create a hierarchical tree. In some cases the dynamics of the network, characterized by the grouping of the clusters, could be approximated from the hierarchical tree. For example, in (Figure 4d,e,f) the oscillator clusters associated with adjacent leaves of the tree fired in a contiguous sequence. The concurrence between the specific coloring and the phase ordering of the clusters depended on the degree of symmetry of the tree. For example, in the complete 9 −partite network simulated in Figure(4), where every node (group of 9 vertices) was equidistant (measured in terms of *J* (*C*)) from all the other nodes, all orderings were possible. All permutations of the nodes of the 9 partite graph were indistinguishable. Different orderings were obtained for different initial conditions (two example sequences with different initial conditions are shown in Figure 5b). In general only some permutations of the nodes are indistinguishable from from each other. If we consider the distance matrix *J* (*C*) as the adjacency matrix of a weighted graph, then the indistinguishable permutations are edge automorphisms of the corresponding graph. permutation of vertices given by matrix *P* is an edge automorphism of this graph if it satisfies *P* ^*T*^ *JP* = *J* or equivalently *J*_*p i,p j*_ = *J*_*i,j*_ *∀ i, j* = 1, 2*, …,* 9 where *p* is a permutation of vertices [16]. For a complete 9 partite graph with 9 vertices in each partition, the non-zero elements of the matrix *J* (*C*) were all 72. There were 8! permutations that did not modify *J* (*C*). For the weighted graph with adjacency matrix *J* (*C*) shown in Figure (4d), there is only one edge automorphism. For the network in Figure(4g) there were 8 possible permutations. The matrix *J* (*C*) contained non-zero elements that were either 18 or 27. Multiple phase orderings of the oscillators were possible as the number of automorphisms of *J* (*C*) increased.

A The clustering observed in the simulations above dependend on the presence of balanced excitation. In fact, in a random 2 partite network, the dynamics could be tuned to go from a state where there was no clustering to a completely synchronous state as this balance was varied figure(2f). Here we examine the role of excitatory-inhibitory balance on the dynamics of the system. In particular, we examine the number of initial conditions that settle to a periodic attractor that can be mapped to a coloring of 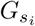. For each value of excitation we simulated the network using 500 random initial conditions. For low values of excitation, very few solutions settled to minimal colorings of the network over the duration of the simulation (figure5a). This was anticipated since we observed similar behavior in the random 2 partite network as well. As excitation increased, an increasing number of solutions settled to periodic attractors. These attractors could be mapped to the colorings of the inhibitory graph 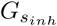 A further increase led to a reduction in the number of periodic solutions. Next, we examined the diversity of solutions. For large values of excitation, we found a preponderance solutions that mapped to colorings with *J* (*C*) matrices that had 18 and 27 as the only non-zero elements (henceforth called 18−27 colorings). We characterized the diversity of solutions as the fraction of 18 − 27 colorings. As the strength of excitation increased this fraction increased monotonically until nearly all solutions were 18 − 27 colorings (Figure 5a). Therefore, when excitation and inhibition were balanced, not only were the periodic attractors more numerous, they were also diverse in terms of the symmetries of the solutions. Why do 18 −27 attractors persist for high values of excitation? We perturbed the system using random independent phase shifts to different neurons. As the size of the perturbation increased, the proportion of solutions that returned to the periodic attractors steadily decreased. Each trace in the figure 5b shows this proportion for 500 instantiations of each periodic attractor. The red lines show the results for the attractors that could be mapped to 18 −27 colorings. These solutions were more stable to perturbations than other colorings. Balanced excitation therefore seems to play a role in stabilizing solutions (colorings) that would otherwise be unstable.

Our results show that a simple network with excitatory-inhibitory balance can encode a large number of spatiotemporal patterns. This presents an interesting opportunity to construct physically realizable systems that can be harnessed to compute. For example, globally coupled arrays of Josephson junctions show (*N* −1)! splay-phase states [17]. As the number of oscillators increase, so do the number of splay-phase states [18]. For neurons, when the network is in a balanced state, we found that the number of attractors was maximal. Thus excitatory-inhibitory balance can potentially lead to attractor crowding [18] causing the system to skip between a large number of periodic attractors. This might explain why balanced excitatory-inhibitory networks show irregular chaotic dynamics [19]. A possible way to generate robust dynamics in neuronal networks is by introducing structural asymmetries through the addition of inhibitory links. These asymmetries reduce the number of possible solutions and stabilize a subset of the remaining solutions [6, 20]. Clues of a Sudoku puzzle can be thought of as constraints that disallow some of the solutions/colorings of the empty Sudoku [11]. Clues may be implemented as additional edges to the Sudoku network that introduce asymmetries in its structure. We can also enforce clues by holding a subset of the oscillators at a fixed phase and allowing the network to evolve under this constraint. A Sudoku grid with enough clues possesses a unique solution. Thus, Sudoku can be fashioned as a content addressable memory system which generates the complete pattern(solution) as output if enough information about the pattern(clues) is given.

## References

[1] G. Laurent, Nature reviews. Neuroscience 3, 884 (2002).

[2] E. Marder and R. L. Calabrese, Physiological reviews 76, 687 (1996).

[3] G. Buzsáki, Neuron 68, 362 (2010).

[4] G. Buzsáki and J. J. Chrobak, Current Opinion in Neurobiology 5, 504 (1995).

[5] M. Rabinovich, A. Volkovskii, P. Lecanda, R. Huerta, H. D. I. Abarbanel, and G. Laurent, Physical Review Letters 87, 068102 (2001).

[6] V. S. Afraimovich, V. P. Zhigulin, and M. I. Rabinovich, Chaos 14, 1123 (2004).

[7] A coloring is a partitioning of the network that assigns vertices that are directly connected to different colors. A partitioning with the fewest number of colors is called a minimal coloring.

[8] C. Assisi, M. Stopfer, and M. Bazhenov, Neuron 69, 373 (2011), arXiv:NIHMS150003.

[9] A. Parihar, N. Shukla, M. Jerry, S. Datta, and A. Raychowdhury, Scientific Reports 7, 1 (2017), arXiv:1609.02079.

[10] R. Lewis, A Guide to Graph Colouring (2016) pp. 1–253.

[11] A. M. Herzberg and M. R. Murty, Notices of the AMS 54, 708 (2007).

[12] R. E. Mirollo and S. H. Strogatz, SIAM Journal on Applied Mathematics 50, 1645 (1990), arXiv:0801.3044.

[13] W. Gerstner, W. Gerstner, W. M. Kistler, and W. M. Kistler, Cambridge University Press (2002) p. 494.

[14] F. S. Neves and M. Timme, 018701, 1 (2012).

[15] R. Diestel, Graph Theory (2005) p. 427, arXiv:arXiv:1102.1087v6.

[16] K. Balasubramanian, Journal of Chemical Information and Computer Sciences 34, 1146 (1994).

[17] S. Nichols and K. Wiesenfeld, 45, 8430 (1992).

[18] K. Wiesenfeld and P. Hadley, Physical Review Letters 62, 1335 (1989).

[19] C. van Vreeswijk and H. Sompolinsky, Science 274, 1724 (1996).

[20] A. S. Pillai and V. K. Jirsa, Neuron 94, 1010 (2017).

